# Gene knock-outs in human CD34+ hematopoietic stem and progenitor cells and in the human immune system of mice

**DOI:** 10.1101/2022.10.06.511235

**Authors:** Daniel A. Kuppers, Jonathan Linton, Sergio Ortiz Espinosa, Kelly M. McKenna, Anthony Rongvaux, Patrick J. Paddison

## Abstract

Human CD34^+^ hematopoietic stem and progenitor cells (HSPCs) are a standard source of cells for clinical HSC transplantations as well as experimental xenotransplantation to generate “humanized mice”. To further extend the range of applications of these humanized mice, we developed a protocol to efficiently edit the genomes of human CD34^+^ HSPCs before transplantation. In the past, manipulating HSPCs has been complicated by the fact that they are inherently difficult to transduce with lentivectors, and rapidly lose their stemness and engraftment potential during *in vitro* culture. However, with optimized nucleofection of sgRNA:Cas9 ribonucleoprotein complexes, we are now able to edit a candidate gene in CD34^+^ HSPCs with almost 100% efficiency, and without affecting their potential for engraftment and multilineage differentiation in mice. The result is a humanized mouse from which we knocked out a gene of interest from their human immune system.

## Introduction

CD34 expression serves as a selective marker of immature hematopoietic cells^1^. In clinical practice, CD34 is used to evaluate and ensure rapid engraftment in HSC transplants^2,3^. CD34+ populations in bone marrow or blood samples are a hematopoietic stem/progenitor mix, of which the majority of cells are progenitors^4^. Human, donor derived CD34+ hematopoietic stem and progenitor cells (HSPCs) populations have become a standard source of cells for allogenic and autologous HSC transplantations^5,6^. As a result, there has been intense interest in genetic manipulating CD34+ HSPCs for use in the treatment of hematopoietic-related diseases, ranging from sickle cell disease (e.g.,^7,8^) to severe combined immunodeficiency^9,10^ to HIV/AIDS^11^ to cancers^12^.

CD34+ HSPCs are also used as *in vitro* and *in vivo* experimental models in conjunction with functional assays (e.g., colony formation, differentiation) and xenotransplantation. For *in vivo* work, novel strains of recipient mice have been developed that better support long term human hematopoiesis and multilineage development of CD34+ cells, including B and T lymphocytes, natural killer cells, monocytes, macrophages and dendritic cells ^13^.

For gene manipulation experiments in HSPCs, there have been numerous attempts to optimize lentiviral (lv) gene expression platforms, including those incorporating CRISPR-Cas9- or RNAi-based elements, using alternate viral envelop proteins, promoters, viral elements etc. (e.g.,^14–17^). However, in our experience such enhanced lv-vectors are still subject one or more biological limitations inherent to HSPCs, either: low transduction efficiencies, transgene silencing, and/or transduction associated toxicity.

Others have begun to develop optimized protocols for gene editing that incorporate delivery of sgRNA:Cas9 nuclease ribonucleoprotein (RNP) complexes to *ex vivo* cultured human CD34+ cells. In some cases, they have reported genomic insertion-deletion (indel) frequencies greater than 90%^18^. However, these protocols still leave room for further improvement. For example, the optimized method reported by Wu. *et al*. requires the use of non-commercially available Cas9, containing an additional NLS^18^. Meanwhile, Modarai *et al*. observed highly variable indel frequencies across patient samples when utilizing the most widely published nucleofection conditions for human CD34+ cells^19^.

Here, we report adapting and optimizing a method of sgRNA:Cas9 RNP nucleofection we use for other primary cell systems^20^ to human CD34+ cells. With this optimized protocol, we consistently achieve a greater than 95% knockout efficiency using standard commercially available reagents and eliminate donor-specific variability in indel frequency previously reported^19^. Utilizing this protocol, we are also able to simultaneously target at least 3 different genes with minimal loss in knockout frequency. We have also optimized the protocol for transplantation of edited human CD34+ cells into humanized mice.

## Results

### Optimization of CD34+ cell nucleofection conditions for delivery of RNA and RNPs

Most publications employing Lonza’s 4D nucleofection technology for delivery of sgRNA:Cas9 RNPs or RNA into CD34+ cells utilize the transfection conditions outlined in the publicly available nucleofection database (knowledge.lonza.com) or very similar conditions. This standard procedure uses Kit P3 and nucleofection program EO-100 with some publications using program ER-100^18,19^. The reported editing efficiencies under these conditions range from 15-90%, with cell viability between 40 and 80% and significant variability in donor to donor editing efficiency^18,19^. Based on unpublished publicly presented data, we decided to try and optimize for a more consistent editing efficiency of CD34+ cells across donors and CD34+ cell source (e.g. bone marrow, G-CSF mobilized) (Figure 1A). Utilizing chemically modified mCherry RNA (Trilink), for quantification of nucleofection efficiency, we compared the standard EO-100 program versus DS-120 and DS-150 (Figure 1B). For both programs we measured a modest increase in nucleofection efficiency versus the standard protocol (EO-100: 83.3%; DS-120: 87.7%; DS-150: 92.7%)(Figure 1C), but a significant increase in levels of mCherry expression within the cells (mCherry high EO-100: 66.3%; DS-120: 79.2%; DS-150: 84%)(Figure 1C). We opted to use the DS-150 program going forward, since it resulted in a larger increase in nucleofection efficiency relative to control conditions (9.4% vs. 4.4%) even though program DS-120 had an overall higher level of mCherry expression. (Figure 1C). We also observe a difference in cell viability between the programs, but our overall cell viability was significantly higher than reported by others (Figure 1C) ^18^ (knowledge.lonza.com).

**Figure 1:**
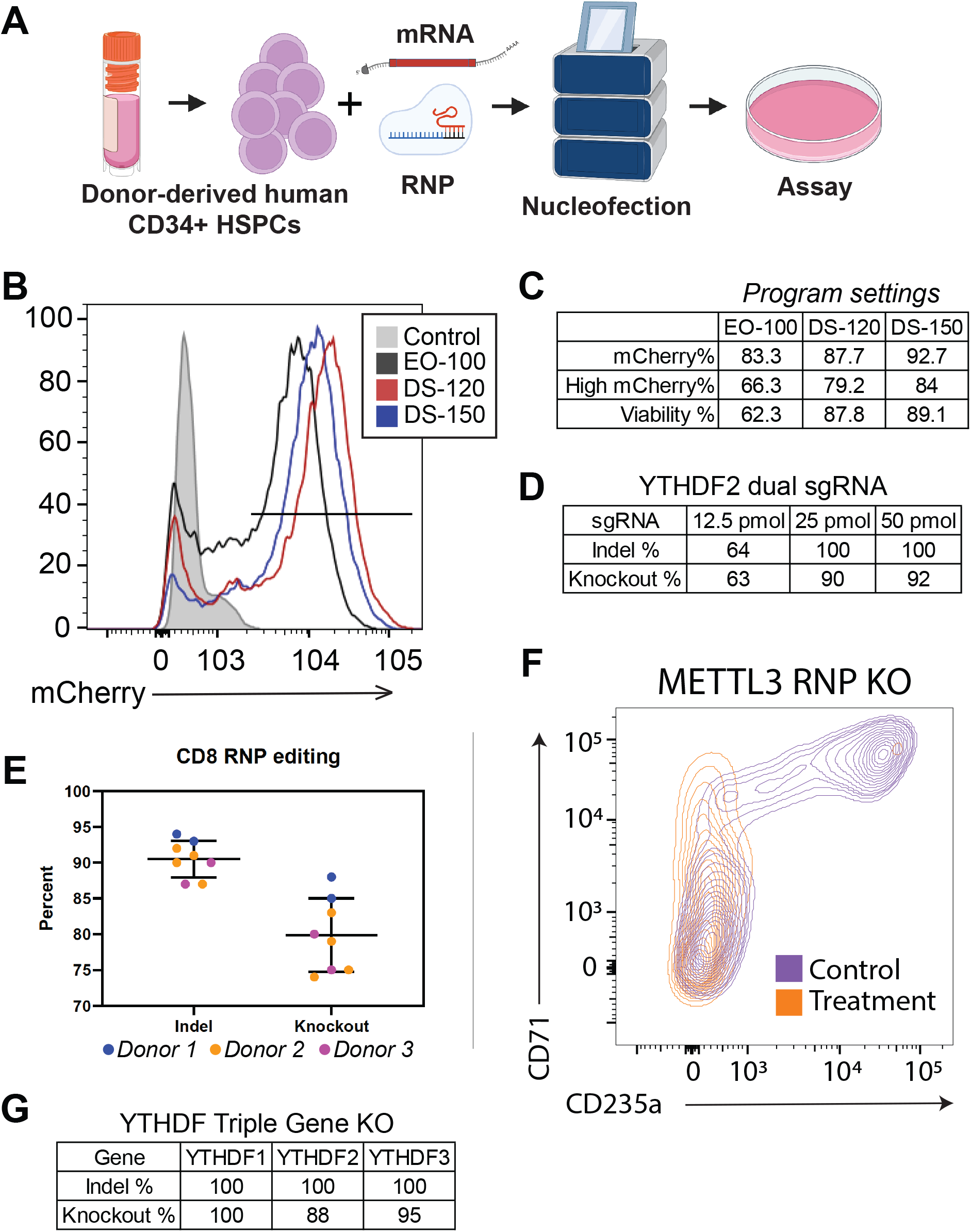
CRISPR/Cas9 RNP editing of human CD34+ hematopoietic stem and progenitor cells. **A**, Overview of sgRNA:Cas9 RNP nucleofection procedure. **B & C**, Optimization of the Lonza nucleofection program selection for CD34+ HSPCs by introducing chemically modified mCherry RNA. Levels of mCherry expression were quantified 2-days post-nucleofection by flow cytometry. **D**, Optimization of input sgRNA amounts when nucleofecting dual sgRNAs for gene knockout. The listed pmole amounts were added for each sgRNA in combination with Cas9 in a 2:1 molar ratio. Three days post-nucleofection gDNA was isolated and CRISPR activity quantified by Sanger sequencing and ICE analysis. **E**, Use of optimized protocol to target CD8 in CD34+ HSPCs from four separate human donors (n=8). Three days post-nucleofection gDNA was isolated and CRISPR activity quantified by Sanger sequencing and ICE analysis. **F**, Representative flow cytometry results comparing the *in vitro* erythroid differentiation phenotype when METTL3 is either KO by CRISPR/Cas9 RNPs or lentiviral shRNA knockdown. Cells were assayed by FACS 7-9 days post-nucleofection or post-transduction. All viable cells from the nucleofection are shown. **G**, Simultaneous triple gene KO by sgRNA:Cas9 RNP nucleofection. CD34+ cells were nucleofected with a pool of RNPs targeting the three YTHDF RNA binding proteins. Each pool contained 25 pmol each of 6 sgRNAs (2 sgRNAs per gene) with Cas9 in a 2:1 ratio. Three days post-nucleofection gDNA was isolated and CRISPR activity quantified by Sanger sequencing and ICE analysis.

Based on previous work in the lab to optimize sgRNA:Cas9 RNP editing in other primary cells and the work of others in CD34+ cells^18^, we were able to use a limited range of sgRNAs amounts and a fixed 2:1 sgRNA to Cas9 ratio for further optimization of sgRNA:Cas9 RNP editing in CD34+ cells. We also standardized on using sgRNAs with 2’-O-methyl 3’ phosphorothioate modifications (in the first and last three nucleotides) from Synthego and 2xNLS SpCas9 from Aldevron. We tested the efficiency of indel formation and gene KO with sgRNA amounts of 12.5, 25, and 50 pmol per sgRNA and 200 thousand cells in a 20 μl nucleofection reaction (Figure 1D) with dual sgRNA targeting YTHDF2. Indel and KO efficiency was quantified as previous described^20^, from Sanger sequencing data and Inference of CRISPR Edits (ICE) analysis provided by Synthego. It should be noted that this is the predicted biallelic indel and KO efficiency.

Maximal gene KO was detected using either 25 or 50 pmol of sgRNA with only a large reduction when using 12.5 pmol (Figure 1D). Interestingly, the reduced editing in the 12.5 pmol sample was due to near complete loss of the fragment deletion between the two sgRNAs. This suggests there is a threshold amount of RNP required to ensure the simultaneous editing necessary for a fragment deletion independent of the activity of the sgRNAs. Given these results, it may be possible to utilize a smaller amount of RNP than we observed if the sgRNAs are pre-validated to have a high KO efficiency independently.

To test consistency of the protocol, we performed sgRNA:Cas9 RNP editing on CD8 using CD34+ HSPCs from three different donors, for a total of eight separate replicates where cohorts of donors were nucleofected on different days (Figure 1E). The results show a remarkable degree of consistency between in the procedure and for these sgRNAs, with predicted indel frequencies averaging >90% regardless of donor, day, or replicate.

### *In vitro* biological validation targeting METTL3 in HSPCs

To validate the optimized RNP nucleofection conditions in a biological context, we opted to target METTL3 for KO. METTL3 is a core component of the N^6^-methyladenosine (m^6^A) mRNA methyltransferase (MTase) complex. m^6^A is an abundant RNA modification that affects the methylated transcripts in a variety of ways including altered stability and translation^21^. This regulation occurs through a host of methyltransferases, demethylases and m^6^A reader proteins, with METTL3 as one of the core components of the methyltransferase complex^22,23^ and we have previously reported that it’s knockdown (KD) by lentivirus delivered shRNAs results in a developmental block to erythropoiesis^24^. Human CD34+ HSPCs expanded for 4 days, as previously described, were nucleofected with RNPs generated from a pool of two METTL3 sgRNAs containing 50 pmol of each. Following the nucleofection the cells were placed in erythroid differentiation conditions^25^. Indel and KO frequency was quantified by ICE analysis in cells collected three days post nucleofection and erythroid differentiation by flow cytometry analysis 6-days post nucleofection (Figure 1F). We detected 91.2% biallelic gene KO and a block to erythroid differentiation consistent with METTL3’s role in erythropoiesis^24^.

### RNP nucleofection for the simultaneous targeting of 3 or more genes

The ability to simultaneously KO multiple genes has utility in a variety of situations. Among these is looking for gene dependencies when trying to identify cancer vulnerabilities ^26,27^ and targeting paralogous gene families where loss of one gene may be functionally compensated for by other genes in the family ^28^. The main technical limitation to target multiple genes by nucleofection is that solutions containing sgRNAs and Cas9 cannot exceed approximately 15-20% of the total volume of the 20 μl reaction. Additionally, very high KO efficiencies must be maintained for all genes, since the more genes that are simultaneously targeted, the greater the effect that small decreases in KO efficiency have on combined targeting efficiency of all the genes. To validate this approach, we simultaneously targeted three YTHDF m6A-binding proteins (DF1, DF2, DF3). Based on our single sgRNA results, we nucleofected 6 sgRNA pools containing 2 sgRNAs per gene and 25 pmol of each sgRNA. The pooled targeting resulted in 100% indel efficiency and 88-100% KO efficiency for all the genes (Figure 1G). When using pre-validated sgRNAs with 95% or greater KO efficiency, this pooled approach should allow the targeting of up to 6 independent genes with approximately 70% of cells being knocked out for all 6 genes and almost all cells having at least a single copy knocked out.

### RNP nucleofected CD34+ cell transplantation into humanized mice results in greater than 95% loss of target gene expression *in vivo*

Previous efforts at gene editing in human CD34+ cells have shown both variability in editing efficiency and persistence of the edits in long-term engrafting HSCs (LT-HSC). Our optimized nucleofection protocol has address the first of these issues. To minimize the loss of LT-HSCs, we further optimized the protocol to minimize the time the cells are cultured *in vitro* before injection into a mouse. In brief (Figure 2A), Human fetal liver-derived CD34+ HSPCs were thawed out and cultured for 4 hours in our standard HSPC expansion media, to allow them to recover from the thawing process. The cells were then nucleofected utilizing our optimized protocol and after a second recovery period of 4 hours, 10,000 cells were injected into the livers of 2-day old preconditioned MISTRG mice as previously described^13^. Initial editing was assessed by ICE analysis after *in vitro* culture of a subset of the cells for 3 days. *In vivo*, KO was assessed by flow cytometry of blood 8-weeks post-transplant.

**Figure 2:**
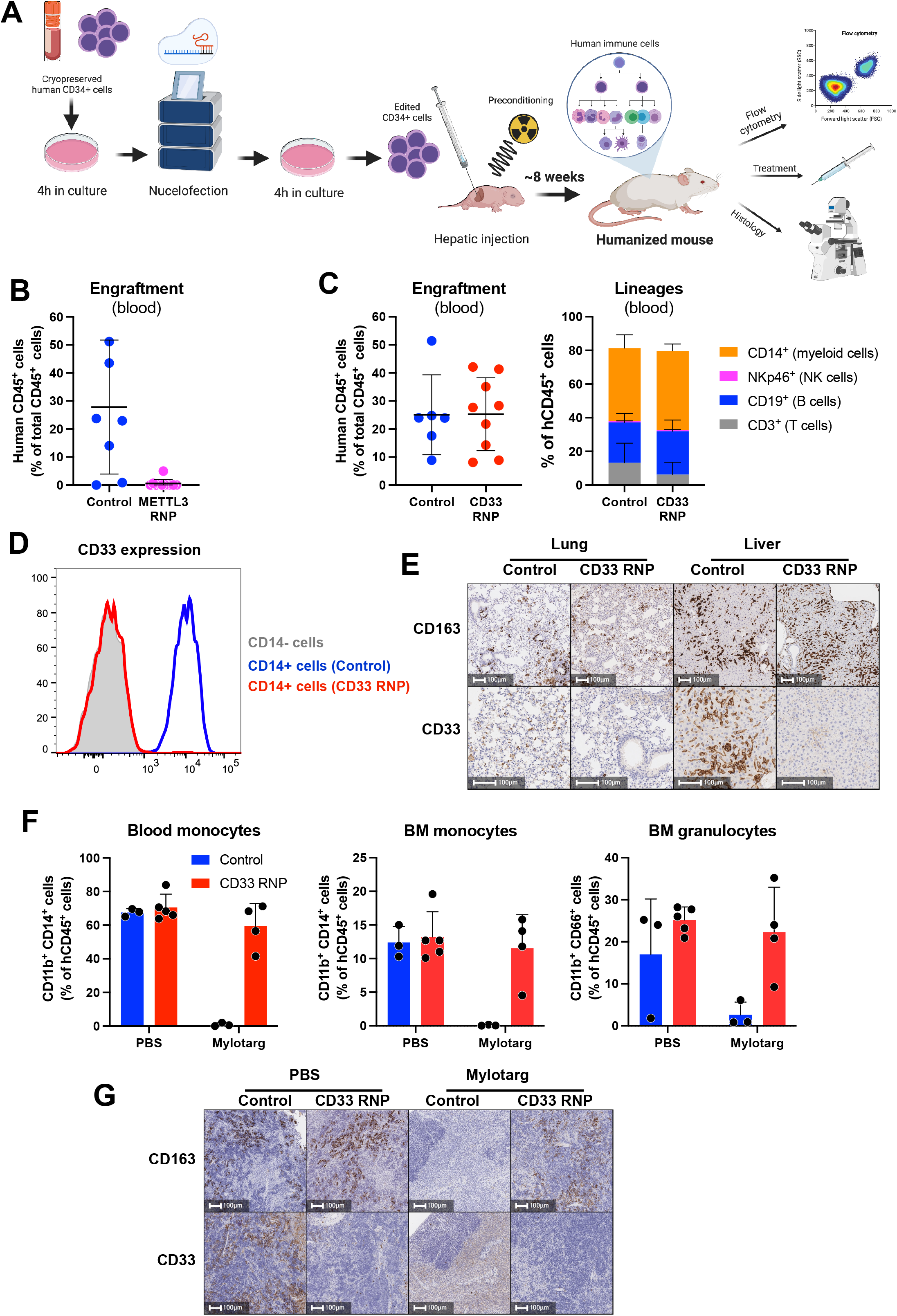
High efficiency in vivo gene KO following transplantation of CRISPR/Cas9 RNP edited human CD34+ cell into the MSTRG humanized mouse model. **A**, The optimized workflow for transplantation of CRISPR/Cas9 RNP edited CD34+ human cells into MSTRG mice. **B**, Frequency of human CD45+ cells in the blood of MISTRG mice 8-weeks after transplantation of fetal CD34+ cells, treated either with control nucleofection or METTL3 RNP. **C**, Frequency hCD45+ cells and lineage differentiation in the in blood of MISTRG mice after transplantation of control or CD33 RNP CD34+ cells. **D**, Expression level of cell surface CD33 by monocytes, identified as Lin-CD14+ cells. **E**, IHC identifying human CD163+ myeloid cells and expression of CD33 in the lung and liver of MISTRG mice. **F**, Frequency of monocytes and granulocytes in the blood or BM of control and CD33 RNP mice, after treatment with Mylotarg. **G**, Identification of CD163+ myeloid cells and expression of CD33 in the spleen of MISTRG mice.

For our first *in vivo* validation study, we again chose to target METTL3. A previous study established that mouse fetal liver HSPCs in which Mettl3 was conditionally deleted have dramatically lower BM engraftment than Mettl3+ cells, owing to the induction of deleterious innate immunity response^29^. To investigate whether this observation is also relevant during human hematopoiesis we utilized the optimized *in vivo* editing and transplantation protocol from above to target METTL3. Our results recapitulate the observation in mice with a near total loss of human hematopoietic cells *in vivo* in the METTL3 KO mice compared to control (CD45+: 0.13% n=11 vs. 27.81% n=8) (Figure 2B). The KO efficiency predicted by ICE analysis was ~90% (not shown).

For our second validation study, we targeted CD33 (SIGLEC3). CD33 is a cell surface receptor for sialic acid proteins primarily expressed on myeloid cells; when KO’d out on its own does not have a developmental phenotype^30,31^. CD33 expression is easily detectable by flow cytometry and immunohistochemistry (IHC), and is the target for the antibody-drug conjugate Mylotarg, which is used to treat acute myeloid leukemia^32^. We found that the nucleofection and targeting procedure did not significantly effect engraftment rates of human CD45+ hematopoietic cells in MISTRG mice, nor their multilineage differentiation (Figure 2C). Remarkably, we observed total ablation of detectable CD33 protein expression from CD14+ myeloid cells by flow cytometry (Figure 2D). Similarly, IHC analysis of lung and liver from CD33 RNP nucleofection abrogated CD33 expression, while the presence of human myeloid cells (identified by the expression of the human macrophage marker CD163) was unaffected (Figure 2E). *In vitro*, we detected approximately 87% KO of CD33 in the fetal liver HSPCs 3-days post nucleofection, suggesting that ICE analysis underestimates Indel/KO formation (not shown).

Further, treatment of mice with Mylotarg showed dramatic clearance of peripheral blood and bone marrow (BM) monocytes (identified by flow cytometry as hCD45+ CD11b+ CD14+ cells) and BM granulocytes (hCD45+ CD11b+ CD66+ cells), which all express CD33 (Figure 2F). However, these cells were entirely resistant to Mylotarg-mediated depletion (Figure 2F). This observation was corroborated by IHC analysis of the spleen, showing that CD163+ marcophages are resisant to Mylotarg in mice transplanted with edited CD34+ cells (Figure 2G). Together, these observations demonstrate that our protocol resulted in the generation of CD33 KO humanized mice.

## Discussion

Here, we describe an improved method for highly penetrant biallelic indel formation in human CD34+ HSPCs via the use programable sgRNA:Cas9 RNP nuclease complexes. This optimized protocol is higher in efficiency than previously published methods^18,19^, including those currently in clinical trials^7^, and works similarly well in adult and fetal HSPCs. Two key advantages of this method are that single or multiple genes can be targeted and that the high efficiency of targeting persists *in vivo* in engrafted cells and their progeny in the BM, peripheral blood, and organ sites. Both of these features allow for integration of this protocol with multiple experimental paradigms, including in combination with clinically relevant agents, such as Mylotarg as we demonstrate.

This method can also be used to create precise deletions using two nearby sgRNA targeting sequences, either in protein coding genes (e.g., removal of exon) or in non-coding DNA elements such as non-coding RNAs and transcriptional enhancers/repressors^33^. Further, given the ability to scale down nucleofections, it is also amenable for small to medium-scale function genetic screens.

However, there are some limitations to this approach. First is expense. The nucleation apparatus used is more expensive than traditional electroporators and has proprietary software and settings (e.g., waveforms, capacitance, etc.) that cannot be readily reversed engineered. The chemically synthesized sgRNAs are also significantly more costly than the DNA oligos used for standard sgRNA cloning. Second, human CD34+ cells are not widely available and do require special handling and culture conditions. Third, because nuclease-based gene editing can result in large chromosomal rearrangements in a portion of cells^34^, multiple controls are warranted for long-term biological assays, especially, *in vivo*. None-the-less, this should be a useful and robust method for gauging gene or genetic element requirement in pre-clinical studies of human HSPCs and their progeny both *in vitro* and *in vivo*.

## Methods

### CD34+ HSPC culture

The G-CSF mobilized CD34+ cells were purchased from the Fred Hutchinson Cancer Center Co-Operative Center for Excellence in Hematology. The cells were quickly thawed in a 37C water bath followed by graduated osmotic equilibration by doubling the total volume with PBS + 0.5% BSA every 2 minutes for a total of 5 doubling. Following centrifugation, the cells were cultured in Stem Span II (Stem Cell Technologies) supplemented with 100 ng/mL each of TPO, IL-6, SCF and FLT-3 ligand. For all the nucleofection optimization experiments the cells were expanded for 3 days prior to nucleofection and cultured in this media for 3 days post nucleofection.

For the erythroid differentiation test the cells were initially expanded as described above. Following nucleofection, the cells were culture as described in Uchida et al. 2018 ^25^, in erythroid differentiation media consisting of IMDM supplemented with 20% Knockout Serum Replacement (Thermo Fisher Scientific) 2 U/mL EPO, 10 ng/mL SCF, 1ng/mL IL-3, 1 μM dexamethasone and 1 μM estradiol for 5 days. The cells were then transitioned to Erythroid maturation media containing in IMDM supplemented with 20% Knockout Serum Replacement, 2 U/mL EPO, 10 ng/mL insulin and 0.5 mg/mL holo-transferrin for 2 days followed by flow cytometry analysis. EPO was purchased from PeproTech and all other cytokines were purchased from Shenandoah Biotechnology.

### Flow cytometry

To monitor erythroid differentiation in the *in vitro* cultures cells were stained with BUV395-CD235a (BD Biosciences 563810) and APC-CD71 (BD Biosciences 341029) and analyzed on a BD Symphony. To evaluate the engraftment levels and multilineage engraftment of human hematopoietic cells in mice, blood was obtained by retro-orbital collection, and RBCs were eliminated by ammonium-chloride-potassium lysis. WBCs were analyzed by flow cytometry, following standard procedures. The following Ab clones were used (all purchased from BioLegend): anti-human Abs CD3 (HIT3a), CD11b (M1/70), CD14 (M5E2 or HCD14), CD19 (HIB19), CD33 (WM53), CD34 (581), CD45 (HI30), CD66b (G10F5) and NKp46 (9E2) and anti-mouse Ab CD45-BV605 (30-F11). Dead cells were excluded by staining with 7-aminoactinomycin D.

### MISTRG mice engraftment studies

De-identified human fetal liver tissues, obtained with informed consent from the donors, were procured by Advanced Bioscience Resources, and their use was determined as non–human subject research by the Fred Hutch Institutional Review Board (6007-827). Fetal livers were cut in small fragments and then treated for 45 min at 37°C with collagenase D (Roche; 100 ng/ml), and a cell suspension was prepared. Hematopoietic cells were enriched by density gradient centrifugation (Lymphocyte Separation Medium; MP Biomedicals), followed by positive immunomagnetic selection with anti-human CD34 microbeads (Miltenyi Biotec). Purity (>90% CD34+ cells) was confirmed by flow cytometry, and cells were frozen at −80°C in FBS containing 10% DMSO.

MISTRG mice (M-CSF^h/h^ IL-3/GM-CSF^h/h^ SIRPα^h/m^ TPO^h/h^ RAG2^−/−^ IL-2Rγ^−/−^) were previously reported ^13,35^. Newborn mice (days 1–3) were sublethally irradiated (150 cGy γ-rays in a [137Cs] irradiator), and ~10,000-12,000 edited CD34+ cells in 20 μl of PBS were injected into the liver with a 22-gauge needle (Hamilton Company), as previously described ^35^. Eight to 10 weeks later, engraftment levels were measured as the percentage of human CD45+ cells among total (mouse and human combined) CD45+ cells in the blood. All animal experiments were approved by the Fred Hutchinson Cancer Research Center (Fred Hutch) Institutional Animal Care and Use Committee (protocol 50941). Mice were treated with two doses (day −4 and day −2) before analysis with PBS or with Mylotarg (gemtuzumab ozogamicin, Pfizer, 5 ug/mouse) administered intravenously by retro-orbital injection.

### Immunohistochemistry

Tissues were fixed in 10% neutral buffered formalin (SigmaAldrich), and embedded in paraffin. Sections were stained with anti-human CD33 (RBT-CD33, Bio SB) or anti-human CD163 (clone EP324, Bio SB) followed by an HRP-conjugated anti-mouse secondary antibody (Leica) and revealed with the peroxidase substrate 3, 3′-diaminobenzidine (Leica).

### CRISPR editing analysis

Nucleofected cells were harvested at indicated timepoints and genomic DNA was extracted (MicroElute Genomic DNA Kit, Omega Bio-Tek). Genomic regions around CRISPR target sites were PCR amplified using Phusion polymerase (Thermo Fisher Scientific) and primers located at least 250 bp outside sgRNA cut sites. After size verification by agaorse gel electrophoresis, PCR products were column-purified (Monarch PCR & DNA Clean-up Kit, New England BioLabs) and submitted for Sanger sequencing (Genewiz) using unique sequencing primers. The resulting trace files for edited cells versus control cells (nucleofected with non-targeting Cas9:sgRNA RNPs) were analyzed for predicted indel composition using the Inference of CRISPR Edits (ICE) web tool ^36^.

## Acknowledgements

We thank members of the Paddison and Rongvaux labs for helpful suggestions, An Tyrrell for administrative support, and the Fred Hutch NIDDK-CCEH Cell Processing Core for providing CD34+ HSPCs. This work was supported by the following grants: NHLBI-U01 HL099993-01, NHLBI-U01 Pilot project HL099997, NIDDK-P30DK 56465-13, NIDDK-U54DK106829, and R01 DK119270-01 (PJP), R01 CA234720 (AR), the Fred Hutchinson Cancer Center Immunotherapy Integrated Research Center (AR and PJP), and the Bezos family (AR). This research was funded in part through the NIH/NCI Cancer Center Support Grant P30 CA015704. We acknowledge Yale University, the University of Zürich, and Regeneron Pharmaceuticals, where MISTRG mice were generated with the support of the Bill and Melinda Gates Foundation. MISTRG mice were developed using Velocigene technology.

## Contributions

D.A.K. and P.J.P. conceived of the initial idea. D.A.K., J.L., P.J.P., and A.R. designed in vivo experiments. D.A.K., J.L., S.O.E. and K.M.M. performed experiments. D.A.K., A.R. and P.J.P. wrote and revised the manuscript.

## References

1. Sidney LE, Branch MJ, Dunphy SE, Dua HS, Hopkinson A. Concise review: evidence for CD34 as a common marker for diverse progenitors. Stem Cells. 2014;32(6):1380–1389.

2. Berardi AC, Wang A, Levine JD, Lopez P, Scadden DT. Functional isolation and characterization of human hematopoietic stem cells. Science. 1995;267(5194):104–108.

3. Berenson RJ, Bensinger WI, Hill RS, et al. Engraftment after infusion of CD34+ marrow cells in patients with breast cancer or neuroblastoma. Blood. 1991;77(8):1717–1722.

4. Majeti R, Park CY, Weissman IL. Identification of a hierarchy of multipotent hematopoietic progenitors in human cord blood. Cell Stem Cell. 2007;1(6):635–645.

5. Chao NJ, Schriber JR, Grimes K, et al. Granulocyte colony-stimulating factor “mobilized” peripheral blood progenitor cells accelerate granulocyte and platelet recovery after high-dose chemotherapy. Blood. 1993;81(8).

6. Holig K, Kramer M, Kroschinsky F, et al. Safety and efficacy of hematopoietic stem cell collection from mobilized peripheral blood in unrelated volunteers: 12 years of single-center experience in 3928 donors. Blood. 2009;114(18):3757–3763.

7. Frangoul H, Altshuler D, Cappellini MD, et al. CRISPR-Cas9 Gene Editing for Sickle Cell Disease and beta-Thalassemia. N Engl J Med. 2021;384(3):252–260.

8. Esrick EB, Lehmann LE, Biffi A, et al. Post-Transcriptional Genetic Silencing of BCL11A to Treat Sickle Cell Disease. N Engl J Med. 2021;384(3):205–215.

9. Rajawat YS, Humbert O, Cook SM, et al. In Vivo Gene Therapy for Canine SCID-X1 Using Cocal-Pseudotyped Lentiviral Vector. Hum Gene Ther. 2021;32(1-2):113–127.

10. Hacein-Bey-Abina S, Pai SY, Gaspar HB, et al. A modified gamma-retrovirus vector for X-linked severe combined immunodeficiency. N Engl J Med. 2014;371(15):1407–1417.

11. Zhen A, Peterson CW, Carrillo MA, et al. Long-term persistence and function of hematopoietic stem cell-derived chimeric antigen receptor T cells in a nonhuman primate model of HIV/AIDS. PLoS Pathog. 2017;13(12):e1006753.

12. Appelbaum FR. Hematopoietic-cell transplantation at 50. N Engl J Med. 2007;357(15):1472–1475.

13. Rongvaux A, Willinger T, Martinek J, et al. Development and function of human innate immune cells in a humanized mouse model. Nat Biotechnol. 2014;32(4):364–372.

14. Dull T, Zufferey R, Kelly M, et al. A third-generation lentivirus vector with a conditional packaging system. J Virol. 1998;72(11):8463–8471.

15. Ramezani A, Hawley TS, Hawley RG. Lentiviral vectors for enhanced gene expression in human hematopoietic cells. Mol Ther. 2000;2(5):458–469.

16. Hong S, Hwang DY, Yoon S, et al. Functional analysis of various promoters in lentiviral vectors at different stages of in vitro differentiation of mouse embryonic stem cells. Mol Ther. 2007;15(9):1630–1639.

17. Amirache F, Levy C, Costa C, et al. Mystery solved: VSV-G-LVs do not allow efficient gene transfer into unstimulated T cells, B cells, and HSCs because they lack the LDL receptor. Blood. 2014;123(9):1422–1424.

18. Wu Y, Zeng J, Roscoe BP, et al. Highly efficient therapeutic gene editing of human hematopoietic stem cells. Nat Med. 2019;25(5):776–783.

19. Modarai SR, Kanda S, Bloh K, Opdenaker LM, Kmiec EB. Precise and error-prone CRISPR-directed gene editing activity in human CD34+ cells varies widely among patient samples. Gene Ther. 2021;28(1-2):105–113.

20. Hoellerbauer P, Kufeld M, Arora S, Wu HJ, Feldman HM, Paddison PJ. A simple and highly efficient method for multi-allelic CRISPR-Cas9 editing in primary cell cultures. Cancer Rep (Hoboken). 2020;3(5):e1269.

21. Wei J, He C. Chromatin and transcriptional regulation by reversible RNA methylation. Curr Opin Cell Biol. 2021;70:109–115.

22. Wang X, Lu Z, Gomez A, et al. N6-methyladenosine-dependent regulation of messenger RNA stability. Nature. 2014;505(7481):117–120.

23. Liu N, Dai Q, Zheng G, He C, Parisien M, Pan T. N(6)-methyladenosine-dependent RNA structural switches regulate RNA-protein interactions. Nature. 2015;518(7540):560–564.

24. Kuppers DA, Arora S, Lim Y, et al. N(6)-methyladenosine mRNA marking promotes selective translation of regulons required for human erythropoiesis. Nat Commun. 2019;10(1):4596.

25. Uchida N, Demirci S, Haro-Mora JJ, et al. Serum-free Erythroid Differentiation for Efficient Genetic Modification and High-Level Adult Hemoglobin Production. Mol Ther Methods Clin Dev. 2018;9:247–256.

26. Meyers RM, Bryan JG, McFarland JM, et al. Computational correction of copy number effect improves specificity of CRISPR-Cas9 essentiality screens in cancer cells. Nat Genet. 2017;49(12):1779–1784.

27. Behan FM, Iorio F, Picco G, et al. Prioritization of cancer therapeutic targets using CRISPR-Cas9 screens. Nature. 2019;568(7753):511–516.

28. Parrish PCR, Thomas JD, Gabel AM, Kamlapurkar S, Bradley RK, Berger AH. Discovery of synthetic lethal and tumor suppressor paralog pairs in the human genome. Cell Rep. 2021;36(9):109597.

29. Gao Y, Vasic R, Song Y, et al. m(6)A Modification Prevents Formation of Endogenous Double-Stranded RNAs and Deleterious Innate Immune Responses during Hematopoietic Development. Immunity. 2020;52(6):1007–1021 e1008.

30. Freeman SD, Kelm S, Barber EK, Crocker PR. Characterization of CD33 as a new member of the sialoadhesin family of cellular interaction molecules. Blood. 1995;85(8):2005–2012.

31. Kim MY, Yu KR, Kenderian SS, et al. Genetic Inactivation of CD33 in Hematopoietic Stem Cells to Enable CAR T Cell Immunotherapy for Acute Myeloid Leukemia. Cell. 2018;173(6):1439–1453 e1419.

32. Petersdorf SH, Kopecky KJ, Slovak M, et al. A phase 3 study of gemtuzumab ozogamicin during induction and postconsolidation therapy in younger patients with acute myeloid leukemia. Blood. 2013;121(24):4854–4860.

33. Hoellerbauer P KM, Arora S, Wu H, Feldman HM, and Paddison P. A simple and highly efficient method for multi-allelic CRISPR-Cas9 editing in primary cell cultures. Cancer Reports. 2020;e1269.

34. Samuelson C, Radtke S, Zhu H, et al. Multiplex CRISPR/Cas9 genome editing in hematopoietic stem cells for fetal hemoglobin reinduction generates chromosomal translocations. Mol Ther Methods Clin Dev. 2021;23:507–523.

35. Voillet V, Berger TR, McKenna KM, et al. An In Vivo Model of Human Macrophages in Metastatic Melanoma. J Immunol. 2022;209(3):606–620.

36. Hsiau T, Conant D, Maures T, et al. Inference of CRISPR Edits from Sanger Trace Data. bioRxiv. 2019.

